# Immobilization in nanocellulose matrix reallocates cyanobacterial proteome resources from growth to bioproduction

**DOI:** 10.64898/2026.06.04.729836

**Authors:** Sara Lupacchini, Emil Sporre, Nashwa L. Attallah, Vishnu Arumughan, Eero Kontturi, Hjalmar Brismar, Thaddeus Maloney, Elton P. Hudson, Valentina Guccini

## Abstract

Solid-state photosynthetic cell factories transform dissolved and atmospheric carbon into chemicals and biomass, offering a platform for the sustainable production of industrially relevant compounds. Yet, how immobilisation affects cell physiology to favour biosynthesis over growth is unclear. Here, we fabricate photosynthetic biohybrids of cellulose nanofiber (CNF) and *Synechocystis* sp. PCC 6803 by osmotic dehydration. Osmotic dehydration resulted in an anisotropic distribution of the cyanobacterial colonies and provided excellent light distribution during cultivation. In an engineered sucrose-secreting *Synechocystis* strain, quantitative proteomics showed that immobilization downregulates ribosomal proteins while upregulating sucrose-synthesis and carbon concentrating machinery, as well as a stress signature of enhanced photosystem II repair, photoprotection, and ROS detoxification. These changes are consistent with a reallocation of constrained proteome resources from growth toward product synthesis. This principle may be extendable to photosynthetic production of compounds beyond sucrose.

## Introduction

Microbial bioproduction enables synthesis of complex chemicals from renewable feedstocks. Photoautotrophic microbes such as cyanobacteria are particularly attractive as they convert inorganic carbon using sunlight and minimal nutrients, often secreting synthesized compounds into the surrounding medium (Hudson, 2021; Liu et al., 2024). In solid-state photosynthetic cell factories, cyanobacteria or microalgae are immobilized within abiotic matrices (Arumughan et al., 2025; Malihan-Yap et al., 2026; Tóth et al., 2024; Yang et al., 2025), where they benefit from more homogeneous light distribution compared to biofilms and suspension cultures, reduce contamination risk, and improved long-term viability (Chua et al., 2024; Jämsä et al., 2018; Rissanen et al., 2021; Yu et al., 2026). Immobilization of cells mechanically constrains expansion and growth, potentially slowing anabolic reactions. In photosynthetic cyanobacteria, this results in an imbalance in the generation and consumption of intracellular energy and reductant. This imbalance triggers a photoprotective response, including detachment of light-harvesting antennae proteins from the photosynthetic reaction centres (Moore et al., 2020). However, an imbalance may be beneficial for bioproduction, as excess energy and reductants could be used for synthesis of a target chemical (Shabestary et al., 2024). Agarose, alginate, and nanocellulose have been extensively investigated as immobilization matrices for cell factories (Lapponi et al., 2022; Sheldon & van Pelt, 2013; Tóth et al., 2024). Alginate matrices have poor mechanical stability which limits long-term use, and low porosity which may limit mass transfer (Kosourov et al., 2017, 2024). Nevertheless, alginate immobilisation has been shown to increase specific production rates of sucrose, succinate, β-phellandrene and ethylene of engineered cyanobacteria relative to planktonic cultures (Tóth et al., 2022; Touloupakis et al., 2016; Vajravel et al., 2020; Valsami et al., 2021). Nanocellulose matrices may have better biocompatibility than alginate, due to higher porosity and greater water retention. Cyanobacteria immobilized in nanocellulose matrices produced hydrogen gas and ethylene over weeks and with high specific productivity (Kosourov et al., 2024; Rissanen et al., 2021). Osmotic dehydration (OsD) as a preparation method can reliably control the porosity of cellulose nanofiber (CNF) hydrogels and hydrated films, achieving a solid content between 0.7 to 12 wt% (Guccini et al., 2022). This control over CNF porosity may overcome diffusion limitations for non-volatile compounds (Levä et al., 2023 Arumughan et al., 2025) and allows study of the relationship between matrix property and cell growth and productivity. Cellular adaptation to immobilisation is a prime determinant of productivity, but there are only few studies of how gene expression is altered upon immobilisation (Sun et al., 2025). Proteomics analysis has detailed the response networks to various stresses in planktonic cell cultures, such as nutrient, oxidative, and osmotic stresses (Haavisto et al., 2024; Jahn et al., 2018; Pérez-Pérez et al., 2006; Plohnke et al., 2015). These studies provide a reference for stresses that may be present in an immobilized system. Furthermore, because proteins are also significant energy investments, and the total protein content in a microbial cell is relatively constrained, changes in proteome composition are interpreted as a re-allocation of cellular resources, particularly in response to changes in nutrient quality (Scott et al., 2014). Immobilisation offers an interesting case, as growth is constrained mechanically and not necessarily by nutrient quality. A reduction in ribosomal proteins could allow a re-allocation of protein resources toward enzymes that synthesize the target chemical. In this work we investigated the cellular response of the cyanobacterium *Synechocystis* sp. PCC 6803 (hereafter *Synechocystis*) immobilized in CNF matrices prepared using OsD, focusing on how the matrix affects cell physiology and chemical production. *Synechocystis* was genetically engineered to synthesize and secrete sucrose. We first characterized the physical properties of pristine CNF hydrated films in terms of optical transparency, porosity and diffusion of reporter molecules. Growth of immobilized *Synechocystis* colonies was monitored by confocal microscopy and chlorophyll content. Sucrose production was evaluated under different light and CO₂ regimes. To investigate the cellular response to immobilization, changes in the proteome were quantified using mass spectrometry. We found that immobilisation in CNFs diverted metabolic flux away from biomass formation toward sucrose secretion in a condition-dependent manner, resulting in higher absolute and specific sucrose production when compared to planktonic culture. Proteomic analysis revealed that growth-associated ribosomes were downregulated, and that proteome space was reallocated to sucrose synthesis enzymes. We propose this reallocation can be exploited to redirect carbon toward other products.

## Result and Discussion

### Physical properties of nanocellulose matrices prepared by osmotic dehydration

An osmotic dehydration (OsD) process was used to make the CNF matrix (**Figure 1**). During dehydration, a CNF suspension (0.2 wt%) and a PEG solution (25 wt%) are separated by a semipermeable membrane (Guccini et al., 2022). The osmotic pressure gradient leads to the dehydration of the CNF suspensions, yielding a homogenous, hydrated CNF film with a 12 wt% solid content. Following dehydration, the films were crosslinked using a CaCl^2^ solution. Hydrated CNF films were optically transparent (**Figure 1B**), and had a spectral transmittance of 70-85% at 500-800 nm and 60% at < 400 nm, similar to values reported for carboxylated CNFs prepared with traditional drop-casting (Fukuzumi et al., 2009; Özkan et al., 2018) (**Figure 1C**). The high degree of the CNFs fibrillation (**Figure S1**) and the formation of a strong interconnected network driven by the osmotic dehydration, including the entangled supramolecular structure and broad size distribution, contribute to the observed optical properties (Kaschuk et al., 2024). The observed value of transmittance is not expected to affect negatively the photosynthetic activity cyanobacteria (Jämsä et al., 2018; Levä et al., 2023; Rissanen et al., 2021). On the contrary, a certain degree of opacity can be beneficial, as light may be scattered by micron-sized features (e.g. entanglements of and bigger nanofibers, aka CNF bundles) within the film’s structure (Kaschuk et al., 2024). Opacity is generally associated with higher haze, which significantly improved cellular growth in immobilised microalgae within agarose hydrogels (Chua et al., 2024). The porosity of the immobilization matrix may affect cell productivity. Previous works compared the growth of *Synechocystis* strains for chemical production in CNF matrices and alginate and found no significant correlation between matrix porosity or composition and the primary metabolic indicators, O_2_ evolution and CO_2_ fixation, during the first 5 days of cultivation (Levä et al., 2023). However, the rate of synthesis of chemicals (ethylene and hydrogen) was consistently higher in nanocellulose than in alginates (Jämsä et al., 2018; Rissanen et al., 2021). The porosity of the CNF hydrated films was assessed using thermoporosimetry (Guccini et al., 2018, 2022), which resolves pores smaller than 200 nm, also identified as mesopores in this paper (Maloney, 2015). This deviates from the official IUPAC definition of mesopores, that have diameter between 2 and 50 nm. The hydrated films consist of 65% macroporous and 35% mesoporous **(Figure 1D)**. The volume of the latter was ca. 3 ml g^-1^, lower than the previously reported value of 5 ml g^-1^ for CNF hydrated films with 12 wt% solid content (Guccini et al., 2022). This discrepancy is attributed to the lower carboxylate content in CNFs prepared here (0.8 mmol g^-1^ compared to 1.5 mmol g^-1^). A lower surface charge results in a lower defibrillation degree and larger nanofibers, decreasing surface area and leading to bigger nanofibers bundles and thus lower porosity (Lu et al., 2017). CNF-based hydrogels have higher porosity than alginates at the same solid weight content (Levä et al., 2023), which is due to differences in morphology of nanosized fibres (CNFs) and polymer chains (alginates), together with differences in swelling capability (Levä et al., 2023; Rissanen et al., 2021). There is no report on correlation between the porosity of the CNF hydrated films and their fluid-gas transport. We measured the fluorescence recovery half-time of fluorescently labelled dextrans (10 kDa, 40 kDa, and 500 kDa) in the CNF matrix using fluorescence recovery after photobleaching (FRAP; **Figure 1E**). FRAP recovery time can be used as a proxy for molecular mobility. For small molecules, mobility is dominated by diffusion, while for large molecules (> 500 kDa), additional effects including network morphology and tortuosity may become relevant (Borges-Vilches et al., 2026). The FRAP recovery time for the 40 kDa probe was 1.6-times longer than the 10 kDa probe, and the FRAP recovery time for the 500 kDa probe recovery time was 2.5 longer, indicating unhindered diffusion. Thus, the CNF network is not expected to limit diffusion of gases to or from the immobilised cyanobacteria or constrain chemical excretion to the media (sucrose MW 0.34 kDa).

**Figure 1:**
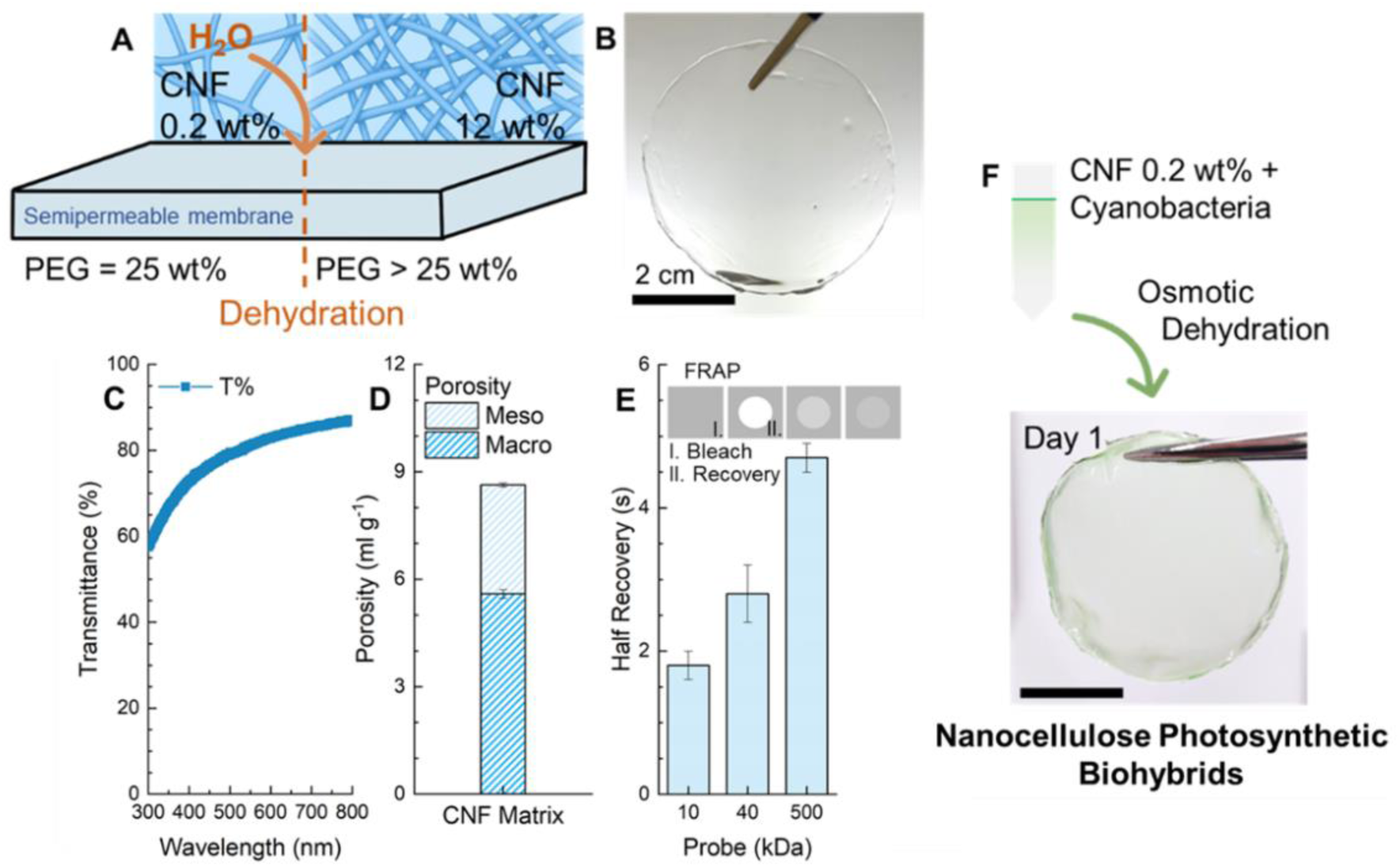
Osmotic dehydration (OsD) and physical chemical properties of the hydrated films. **(A)** representation of densification of the cellulose nanofiber (CNF) network from the suspension (0.2 wt%) to the hydrated films (12 wt%) induced by OsD; **(B)** picture of the CNF hydrated films obtained by OsD; **(C)** transmittance, **(D)** porosity measured by thermoporosimetry and **(E)** fluorescence half recovery after photobleaching (FRAP) of the hydrated films; **(F)** Photosynthetic biohybrids prepared with CNF (0.2 wt%) and *Synechocystis* (scale bar 2 cm).

### Cyanobacteria grow uniformly in the CNF matrix

*Synechocystis* cultures were immobilized in the CNF matrix to obtain the photosynthetic biohybrid. The spatial distribution and growth of the cell colonies in the matrix were characterised with confocal microscopy. **Figure 2A** shows the CNF cross section (thickness 200 μm) after 1 day of incubation under low-intensity illumination (3 µE). The *Synechocystis* colonies appear as light gray features immobilized within the dark CNF matrix as discrete spherical colonies and distributed anisotropically along the CNF matrix cross-section (**Figure 2A and S2**). Most cells are within 100 μm from the top surface facing the light during cultivation, which is due to the OsD preparation method and not cell migration. During preparation, water flows from CNF-cell suspension across the semipermeable membrane which increases the solid content and results in a higher number of cells near the membrane surface. An enrichment of cells near the surface (100 μm), contrasts with the uniform entrapment in alginate beads and the homogeneous distribution in thin other carboxylated CNF- or alginate-based hydrogels (Jämsä et al., 2018; Rissanen et al., 2021). Imaging of colony growth within the top 15 μm showed that the cells proliferate from the initial inoculation point as compact, roughly spherical colonies (**Figure 2B**). As colonies expand, the immobilisation matrix must accommodate the cell proliferation; CNF matrices are particularly advantageous compared to other polymer matrices because their fibrillar network can better tolerate dilatation without losing structural integrity, imparting less mechanical friction and shear on the cells. The rate of colony growth was uniform within 15-100 μm from the film surface over 14 days (**Figure 2C, Figure S3)** indicating a uniform microenvironment. The average colony dimension at the bottom of the film was larger compared to those at the top (**Figure 2D**). This difference could be due to hydration, as the lower regions of the film likely retain moisture more effectively, providing a more favourable microenvironment for cell growth. After 14 days, colonies near the top surface became larger (**Figure S4**), suggesting that light availability was limited at the bottom of the film. It is notable that self-shading effects in the bio-hybrid films become apparent only after day 14. A combination of optical and microstructural properties and cell seeding density may minimize internal gradients during early growth stages and delay the onset of light attenuation effects. We measured the bulk cell growth and cell photosynthetic efficiency of *Synechocystis-*CNF biohybrids in three conditions (**Table 1**); very low light/low CO_2_ (VLL), low light/low CO_2_ (LL), and high light/high CO_2_ (HH). Films were in a bath of growth media in a standing condition without agitation. The cell growth rate in the CNF matrix increased with light intensity (**Figure 2E**). The photosynthetic quantum yield Y(II) of *Synechocystis* was similar in planktonic culture before and immediately after immobilisation in CNF matrix, which contrasts with the drop in photosynthetic efficiency reported when using conventional solvent casting approaches (Jämsä et al., 2018; Rissanen et al., 2021). During growth, Y(II) declined only slightly under VLL and LL conditions, while under HH condition it was lower and decreased more rapidly, consistent with the observed reduction in Y(II) at faster growth rates in planktonic culture (Touloupakis et al., 2015). Although 50 µE is not considered a high-light condition for *Synechocystis*, cell immobilisation can promote an over reduced photosynthetic electron transport chain, leading to ROS generation and oxidative stress (Nikkanen et al., 2021; Sun et al., 2025; Vajravel et al., 2025).

**Figure 2:**
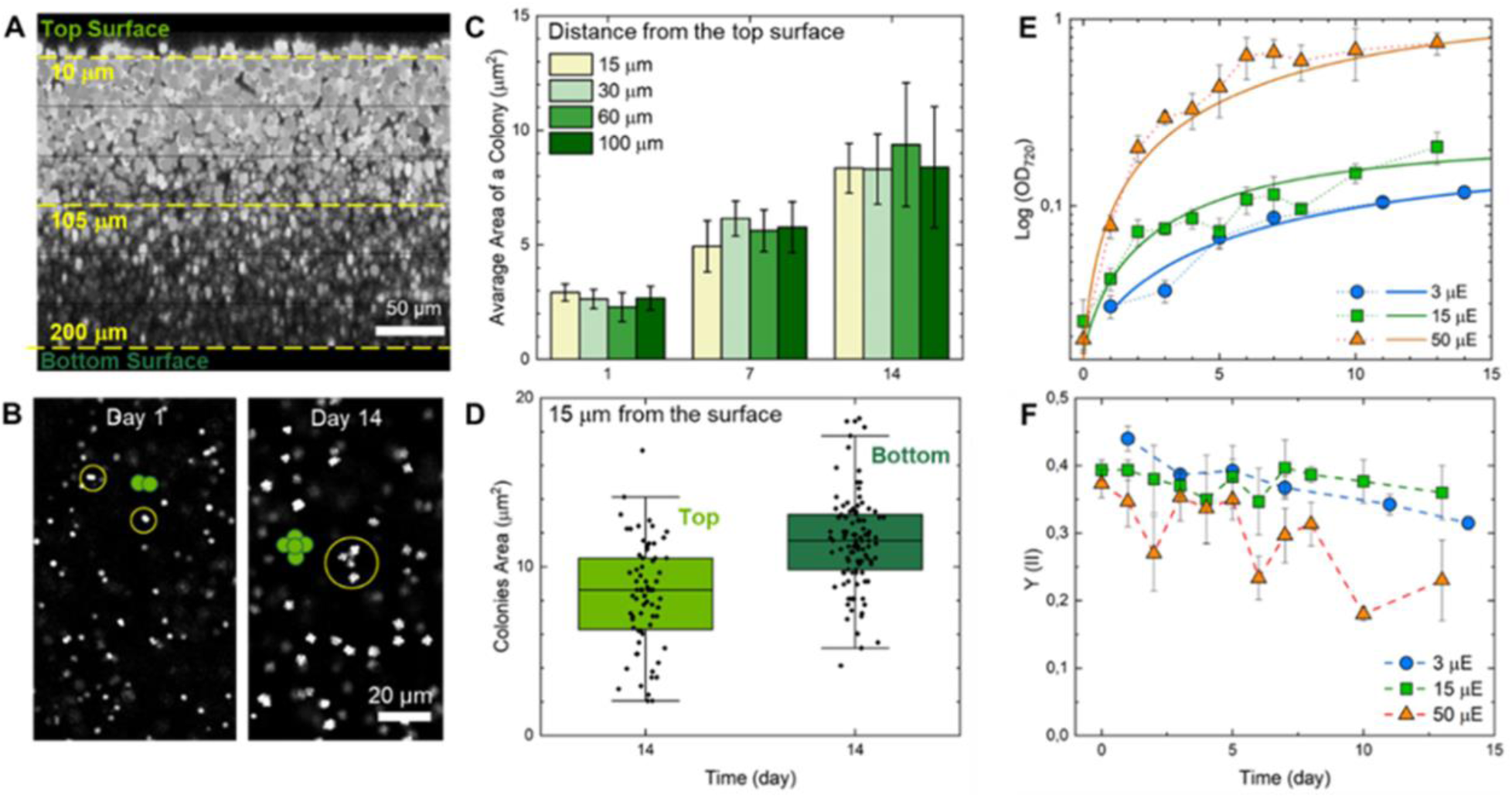
Morphological characterization and growth at different conditions of the immobilized cyanobacteria *Synechocystis.* **(A)** Cross-section of the CNF photosynthetic biohybrids showing the anisotropic distribution of the photosynthetic cells; **(B)** microbial colonies imaged by confocal microscopy at 15 μm depth from the biohybrid top surface (directly exposed to the light); **(C)** quantification of the dimension of the microbial colonies at difference depth; **(D)** average dimension of the colonies at 15 μm depth from the top and bottom surface (in contact with the Petri dish); **(E)** Optical density at 720 nm (OD_720_) and **(F)** quantum yield (YII) of the photosynthetic biohybrids at different condition (3 or 15 μE, 0.04% CO_2_ and 50 μE and 1% CO_2_ enriched).

**Table 1.** Cultivation conditions for CNF immobilized and planktonic *Synechocystis*.

| Condition | Light intensity<br>($\mu\text{mol m}^{-2} \text{s}^{-1}$) | CO<sub>2</sub> in gas phase<br>(%) |
| --- | --- | --- |
| Very low light/Low CO<sub>2</sub><br>(VLL) | 3 | 0.04 |
| Low light/Low CO<sub>2</sub><br>(LL) | 15 | 0.04 |
| High light/High CO<sub>2</sub><br>(HH) | 50 | 1.0 |

### Sucrose synthesis by *Synechocystis* is significantly enhanced by immobilisation in CNF matrices

To test the capability of CNF to promote bioproduction, we immobilized a *Synechocystis* strain genetically engineered to secrete sucrose (SLS21). Key genetic modifications include deletion of *ggpS* to eliminate the competing glucosylglycerol sink, deletion of invertase-*sll0626* to prevent intracellular sucrose catabolism, and plasmid-based overexpression of sucrose phosphate synthase (*sps*) and a heterologous *E. coli* sucrose permease *cscB,* both driven by a strong constitutive promoter P*cpc560*. Together, these changes enabled high sucrose secretion under non-saline conditions (**Figure S5**) (Ducat et al., 2012; Kirsch et al., 2018, 2019; Santos-Merino et al., 2023; Thiel et al., 2018). SLS21 was immobilised in CNF matrix (hydrated film disc in **Figure 3A, B**) in the LL and HH conditions. Both conditions were also tested with vessel shaking, to assess how mass transfer from bulk liquid into the CNF influences cell growth and metabolic partitioning of fixed carbon. In a preliminary experiment, photosynthetic yield Y(II) was stable at 0.3 in both LL and HH non-shaking conditions as cells grew in the matrix until 14 days. At 19 days, Y(II) declined to 0.2 in both conditions (**Figure S6)**. This trend likely reflects a diffusion limitation in the static state, where reduced gas exchange and slower replenishment of dissolved inorganic carbon negatively affect photosynthetic efficiency. Sucrose adsorption to the CNF matrix was ruled out by quartz crystal microbalance with dissipation (QCM-D) measurements using a CNF thin film as a model surface, which showed negligible frequency and dissipation shifts upon sucrose addition, confirming that sucrose transport through the matrix is not impeded by interfacial interactions (**Figure S7**). Representative photographs of SLS21 embedded in hydrated films discs at day 6 illustrate visible differences in biomass accumulation across the four conditions (**Figure 3A, B**).

**Figure 3.**
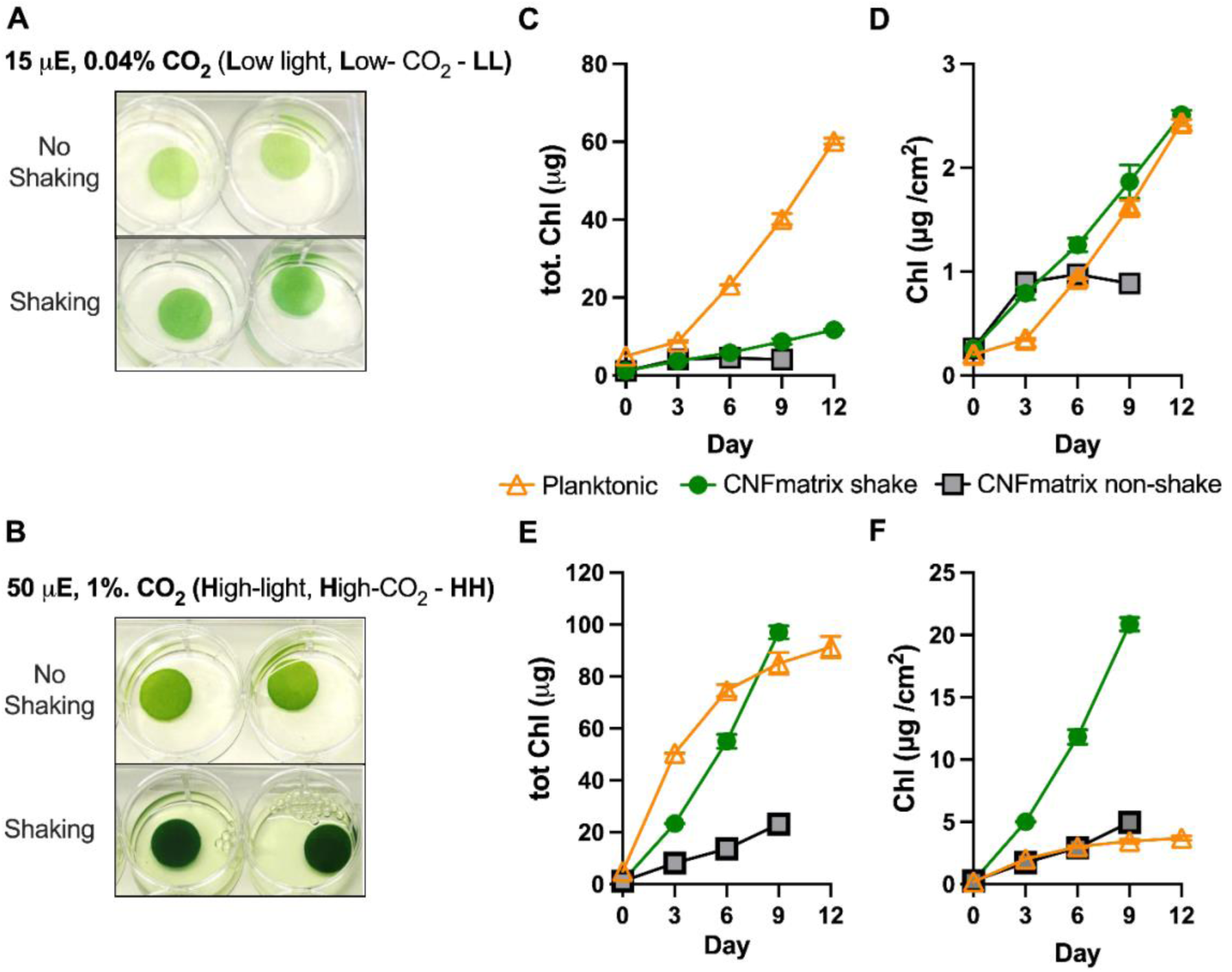
Cell growth determined by chlorophyll content in immobilised and planktonic SLS21. Representative photographs comparing CNF-embedded SLS21 at day six of cultivation in BG11+ 0.4 M NaCl in the four conditions tested: (**A-B**) 15 µE, ambient CO₂, (LL); non-shaking and shaking; 50 µE, 1% CO₂ (HH) non-shaking shaking (**C-D**) Total chlorophyll content (μg) during cultivation under (**C**) 15 µE ambient CO₂ and (**D**) 50 µE with 1% CO₂. (**E-F**) Chl concentration normalised by the area of the CNF matrix disc or of the well plate (μg mL^-1^ cm^-2^).

Cell growth was measured using chlorophyll as a proxy for biomass. CNF cultures showed poor growth in the LL no-shaking condition (total chlorophyll 4 µg at day 9), which was overcome in the LL shaking condition (total chlorophyll 12 µg). Planktonic cultures in the LL condition reached 60 µg. The higher total growth in planktonic culture is partly due to a larger illuminated surface area of the well plate (25 cm²) relative to the CNF disc (5 cm²) (**Figure 3C**). When normalised to illuminated surface area, the CNF and planktonic chlorophyll accumulation were similar ∼ 2 µg Chl cm⁻² at 9 days, while CNF non-shaking cultures plateaued at ∼ 1 µg Chl cm⁻² (**Figure 3D**). In the HH condition, CNF shaking and planktonic cultures accumulated comparable total chlorophyll (∼ 97 and ∼ 91 µg after 9 days, respectively), both substantially exceeding CNF non-shaking cultures (∼23 µg at day 9), confirming that mass transfer of gas or nutrients is a primary growth-limiting factor regardless of external light and CO₂ supply (**Figure 3E**). Notably, when normalised per unit area, the CNF shaking cultures reached 20 µg Chl cm⁻² at day 9, approximately four times higher than planktonic (∼ 4 µg Chl cm⁻²), demonstrating that the CNF matrix concentrates biomass more efficiently per unit illuminated area when mass transfer is ensured (**Figure 3F**). We measured sucrose production by quantifying sucrose in the surrounding culture medium over time. In LL condition, planktonic cultures and CNF-shaking cultures produced comparable sucrose concentrations, reaching 1800 and 1660 µg mL⁻¹ respectively after 12 days. This result is remarkable as planktonic cultures accumulated approximately 4-times more total biomass under the same conditions (**Figure 3C**). The CNF no-shaking culture produced significantly less sucrose (∼ 400 µg mL⁻¹) (**Figure 4A**), consistent with the growth restriction. In the HH condition, CNF-shaking culture showed the highest absolute sucrose accumulation, ∼ 2800 µg mL⁻¹ at 9 days, compared to ∼ 1700 µg mL⁻¹ for planktonic and ∼ 1600 µg mL⁻¹ for CNF-non-shaking (**Figure 4B)**. The CNF immobilized SLS21 in HH condition had among the highest sucrose productivities (39.5 mg L^-1^ h^-1^) reported for *Synechocystis* (Domínguez-Lobo et al., 2025; Kirsch et al., 2018; Santos-Merino et al., 2023; Thiel et al., 2019; Tóth et al., 2022). Normalization of sucrose production to total chlorophyll content revealed a prominent advantage of immobilization, particularly under LL conditions where CNF-shaking cultures showed 4 times higher specific sucrose production than planktonic cultures (**Figure 4C**). In the HH condition, the specific sucrose production was not higher than planktonic, due to higher cell growth in this condition (**Figure 4D**). These results suggests that immobilisation can redirect fixed carbon away from biomass accumulation and toward sucrose secretion, consistent with previous observations in alginate-immobilised sucrose-secreting *Synechocystis* (Tóth et al., 2022). Additionally, cultivation conditions such as light and CO_2_ levels, determine the metabolic regime of CNF immobilized cells: LL promotes a low-growth, sucrose synthesis state with high per-biomass sucrose yield, while HH favours higher growth at the expense of sucrose synthesis.

**Figure 4.**
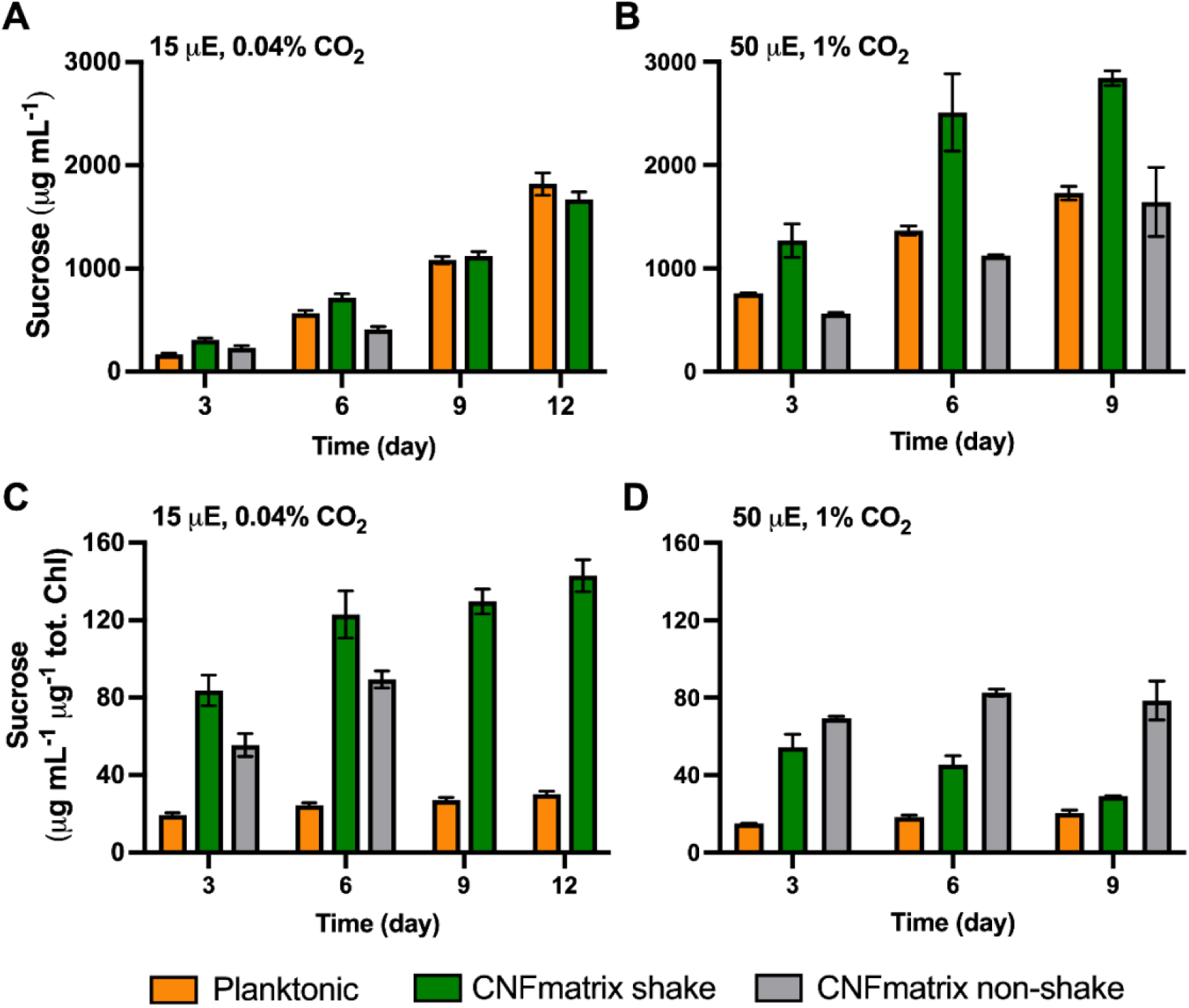
Study of sucrose-producing SLS21 in planktonic cultures and CNF matrix shaking and non-shaking conditions. Total sucrose production (μg mL⁻¹) measured in the extracellular media (**A**) under LL (LL-15 µE light, 0.04% CO₂) and (**B**) (HH-50 µE, 1% CO₂). Sucrose yield normalized to chlorophyll (μg sucrose per μg Chl) (**C**) under LL (LL-15 µE light, 0.04% CO₂) and (**D**) (HH-50 µE, 1% CO₂) Data points are mean ± SD (n = 3).

### Immobilization of cyanobacteria re-allocates proteome from growth to product synthesis

To investigate the trade-off between cell growth and sucrose synthesis, we compared the proteomes of CNF immobilized cells and planktonic cells grown in the LL condition using LC/MS at 3, 6 and 9 days after cultivation start, where >1500 proteins were detected in all samples. Below, we present proteome differences at 9 days, the full dataset is in **Supplemental Data**. Comparison of these revealed significant differences across energy and carbon metabolism, consistent with our hypothesis that CNF immobilization imposes mechanical and physiological stress that constrains growth and diverts carbon metabolism from biomass to sucrose production (Moore et al., 2020; Sun et al., 2025; Tóth et al., 2022) (**Figure 5, Figure S8, Figure S9**). Photosystem II (PSII) core subunits and proteins associated with PSII repair and maintenance were significantly increased in CNF immobilized cells, while the extrinsic luminal proteins PsbO, PsbU and PsbV were strongly decreased (**Figure 5A and 5B**, detailed representation in **Figure S8)**. These three proteins stabilize the Mn_4_CaO_5_ cluster in the oxygen-evolving complex (OEC), which provides electrons that drive photosynthesis (Bricker et al., 2012). Plastocyanin (PetE), which shuttles electrons from Cyt b6f to PSI, was also decreased, indicating down-regulation of linear electron flow (Ivanov et al., 2012). FtsH which degrades damaged PSII subunits during repair cycle, was also increased (Krynická et al., 2015). This expression pattern matches previously reported proteomics changes in *Synechocystis* planktonic cultures when PSII is under photodamage (Nishiyama et al., 2001; Theis & Schroda, 2016). Immobilized cells also upregulated photoprotective mechanisms, including the orange carotenoid protein (OCP), which dissipates excess light energy absorbed by the phycobilisome antenna, and photosystem I-ferredoxin-flavodiiron proteins which together dissipate excess electrons from photosynthetic electron transport that cannot be used for NADPH generation and carbon fixation (Miller et al., 2022; Zhang et al., 2009) (**Fig 5A and Figure S8**). Overall, proteomic changes in photosynthesis proteins resemble the high-light photoinhibition response of *Synechocystis* planktonic cultures (> 800 µE) (Cordara et al., 2018), despite the low incident light used here (15 µE). This suggests an imbalance between photosynthetic electron supply and the limited downstream sinks available when cell division is constrained, which can lead to backpressure in the photosynthetic electron chain and generation of ROS. Supporting this interpretation, ROS-detoxification enzymes catalase-peroxidase (KatG), superoxide dismutase (sodB), peroxiredoxins (Prx5) were upregulated in CNF-immobilized cells (**Figure 5B**). These results are also consistent with a recent study that measured increased ROS levels in agarose-immobilized *Synechococcus* cultures using fluorescent probes, as well as increases in expression of genes related to ROS stress (Sun et al., 2025). Immobilised SLS21 cells showed reduced growth rate but maintained and even enhanced sucrose biosynthesis and export (**Figure 3**, **Figure 4**). The abundance of ribosome proteins, a proxy for cell growth rate, was markedly reduced (**Figure 5B**, violin plot). Ribosome proteins account for up to 10% of the total proteome mass, even at low growth rates, and reduction of these is typically in coordination with increases of other metabolic enzymes (Jahn et al 2018). Indeed, the sucrose-synthesising enzyme SPS and the heterologous sucrose transporter CscB both increased by 100% (log_2_fold-change in abundance of 1 and 1.03 respectively, **Figure 5B-C**). The core components of the carbon concentrating mechanism (CCM) were also highly upregulated, including carbonate transporters BicA and CmpA and the NDH-1 carbonic anhydrase subunit CupA (**Figure 5C, Figure S9).** These are the primary bicarbonate uptake transporters in cyanobacteria, and their increase suggest that immobilized SLS21 may perceive a carbon limitation compared to planktonic cells, either due to poor transport of bicarbonate to or within the matrix, or as an indirect consequence of mechanical constraint on growth. Several TCA cycle-associated enzymes, including elements of the GABA shunt significantly increased in abundance (**Figure 5C, Figure S9).** The TCA cycle in *Synechocystis* has been shown to carry only minor flux under autotrophic conditions (Young et al., 2011), so these changes are more likely to support redox poise and anaplerosis than high respiratory activity. The upregulation of malate dehydrogenase (MDH, log2FC 3.49) and malic enzyme (ME, log2FC: 1.06) highlight an elevated malate pool (Figure S9). MDH preferentially catalyses the reductive reaction (oxaloacetate to malate) and it has been associated with oxidative stress in *E. coli*, while ME is the primary enzyme oxidising malate to pyruvate and CO_2_ with NADPH regeneration. Their co-upregulation suggests a possible redox rebalancing mechanism (Katayama et al., 2022; Takeya et al., 2018).

**Figure 5:**
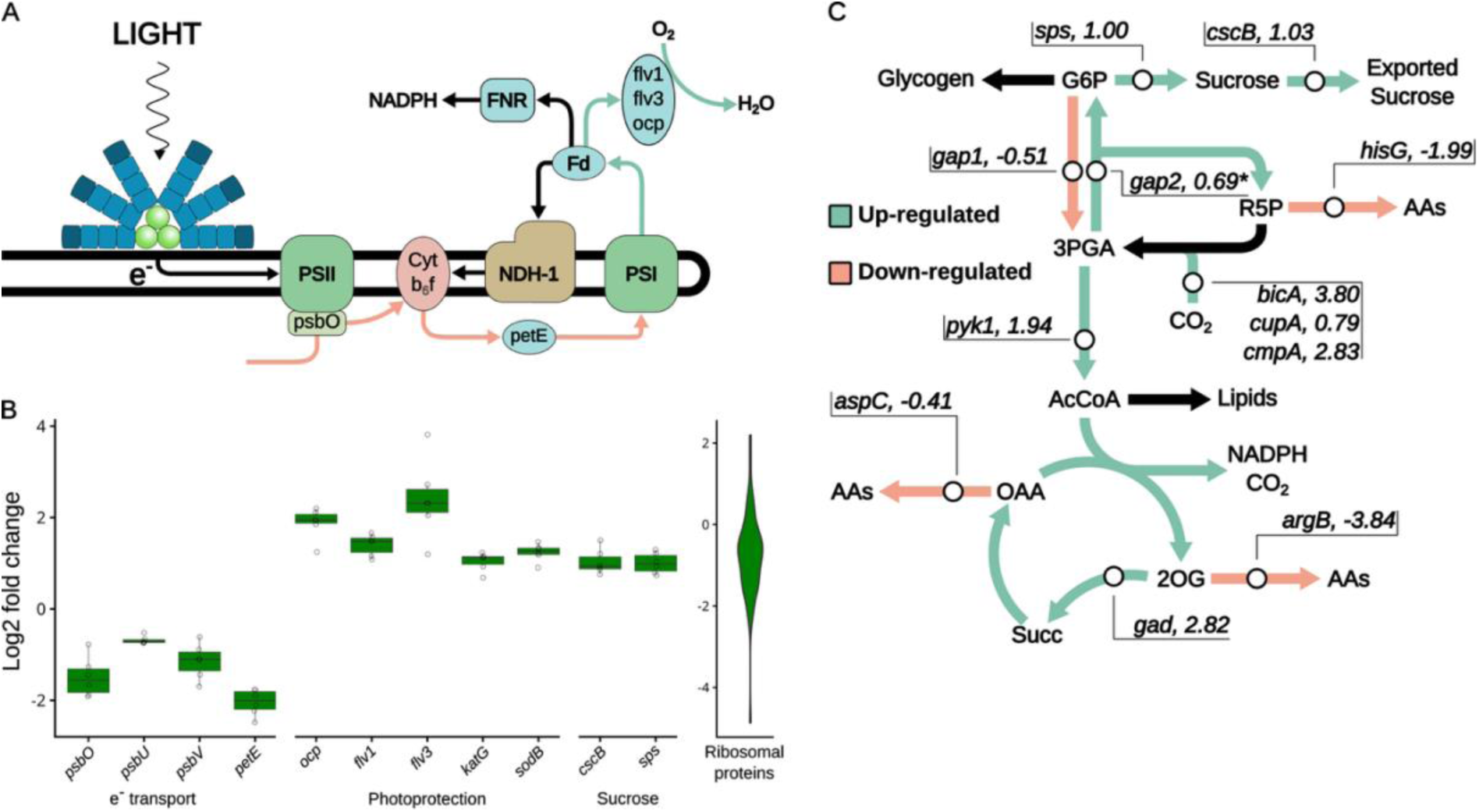
Schematic overview of changes in the proteome of CNF-immobilised, sucrose-secreting *Synechocystis* SLS21 grown under LL shaking condition (15 µE light, 0.04% CO₂) as compared to planktonic cultures at 9 days. (**A**) Overview of photosynthetic electron transport chain where green arrows indicate positive abundance changes, and red indicate negative abundance changes. See **Figure S8** for a more complete overview. (**B)** Changes in abundance of selected proteins in electron transfer, photoprotection, sucrose synthesis, and ribosomal proteins. All changes were statistically significant with adjusted p-value < 0.05 (**C)** Overview of *Synechocystis* central carbon metabolism with selected proteins highlighted and log2 fold-change in abundances in immobilised cultures compared to planktonic. All changes were statistically significant with adjusted p-value < 0.05 except *gap2 adjusted p-value = 0.07. See **Figure S9** for a more complete overview.

## Conclusion

We present CNFs prepared by osmotic dehydration (OsD) as a matrix for immobilisation of cyanobacteria cell factories. CNF matrices prepared by OsD have optimal transparency in the photosynthetic spectrum, a predominantly macroporous architecture, and unrestricted diffusion of dextrans of up to at least 500 kDa as measured by FRAP. Osmotic dehydration drives the enrichment of cyanobacterial colonies within the top 100 μm of the films. We exploited this anisotropic distribution by orienting the cell-dense layer toward the incident light to maximize photon capture from the onset of cultivation. Using an engineered sucrose-secreting strain, we showed that light, CO₂ availability, and mass transfer jointly determine whether immobilized cells operate in a growth-dominated or production-dominated regime. Under low light with adequate mass transfer, immobilisation redirected fixed carbon away from biomass to sucrose synthesis, yielding up to 4-times higher specific productivity than planktonic cultures and absolute titers among the highest reported. Proteomics analysis provided a detail of this shift and points to a broader principle that mechanical growth restriction reallocates the cellular proteome. Ribosomal proteins were downregulated while sucrose synthesis and the carbon concentrating mechanism were upregulated, consistent with a constrained proteome reallocating from growth to sucrose synthesis. This reallocation was accompanied by a stress response that included PSII repair, photodamage, and ROS detoxification, indicating that immobilised cells experience an imbalance between photosynthetic electron supply and downstream sinks. Together, these results show that immobilization is not just a containment strategy but can be a physiological lever for photosynthetic bioproduction. The proteome reallocation mechanism should be transferable to products beyond sucrose but requires validation of the reallocation model. Strengthening PSII repair and detoxification may prolong cell metabolic activity when embedded in matrices.

## Methods

### Preparation of the hydrate CNF films and photosynthetic biohybrids by osmotic dehydration

The carboxylated CNFs were prepared and characterized following previous protocols (Pääkkönen et al., 2015) (details are summarized in the Supporting Information) and used to prepare the CNF hydrated films by osmotic dehydration (OsD) setup (Guccini et al., 2022) (Figure S1). Prior to contact with cyanobacterial cultures, all materials were sterilized under UV light for at least 30 min. A stock suspension of CNF (1 wt%) was diluted to 0.3 wt% and homogenized at 12,000 rpm using an Ultra-Turrax. In parallel, liquid cultures with an OD_730_ of 1-1.5 were centrifuged at 3500 rpm for 10 min at room temperature, washed once with fresh BG11 medium, and resuspended to a final OD_730_ of 2.5. For the immobilization, 20 g of the diluted CNF was mixed with 2.8 mL of the prepared photosynthetic culture in a 50 mL falcon tube. The mixture was vortexed to obtain a homogeneous dispersion, and entrapped air bubbles were removed in vacuum chamber. An analogous OsD setup described by Guccini et al., was used (Guccini et al., 2022). The culture-CNF suspension was poured into the inner cup, which was then placed inside the polyethene glycol (PEG)-containing outer cup (PEG molecular weight of 35 kDa), ensuring membrane (6-8 kDa cut-off) contact with the PEG solution. Osmotic dehydration proceeded overnight on a magnetic stirrer at room temperature, after which the so obtained hydrated films with immobilised photosynthetic culture was crosslinked with a 2 wt% CaCl₂ solution. After this the photosynthetic biohybrid was washed with Milli-Q water and resuspended in BG11 medium. The resulting biohybrid was cut either into rectangular strips (approx. 1 × 3 cm) for measuring the optical density and quantum yield measurements using an AquaPen AP-C 100 hand-held fluorometer (Photon Systems Instruments, Czech Republic), or into circular discs with outer diameter ∼ 17 mm and height 200 μm to measure the sucrose production by assay and to carry out the proteomic analyses.

### Characterisation of the CNF hydrated films

The optical transparency of the pristine hydrated films was measured in terms of transmittance (%) in the wavelength interval between 300 and 800 nm (Guccini et al., 2022). A strip of the hydrated films was cut to fit in the optical transparence cuvette containing deionized water (which served as a blank). The measurements were done using a Shimadzu UV-2550 VMT1 spectrometer. The porosity was measured by thermoporosimetry. These measurements were done in triplicates using a differential scanning calorimeter (DSC), model Mettler Toledo DSC3+ (Mettler-Toledo Intl. Inc. Instrument, USA) equipped with an intracooler. Briefly, the films were blotted with paper to remove the excess water and hermetically sealed in a 40 μL aluminium pan. Both the mass of the hydrated films and the pan were recorded. The combined mass was measured before and after the DSC measurement to make sure that the seal was maintained throughout the measurements. First, the temperature was lowered to - 50°C at a rate of 20 K min^-1^, during which the freezable water in the hydrated films crystallized. Second, the temperature was increased to -0.2°C until the melting transition was completed (corresponding to the melting of the confined water), to prevent supercooling during the subsequent recrystallisation step. 0.2°C is the upper limit of the DSC measurement, before bulk water begins to melt. It corresponds to a Gibb-Tomson pore diameter of 200 nm. The temperature was then decreased at a rate of 2 K min^-1^ to - 50°C, resulting in an exothermic peak from which the freezing bound water (FBW) was calculated, corresponding to the mesoporosity. This calculation was done using a modified version of the Gibbs-Thomson Maloney (Maloney, 2015). The macroporosity was calculated assuming that all the free water was located outside the mesopores and that no loosely absorbed water was present at the surface of the films. This was achieved by appropriately blotting the films before the measurement. The fluid transport of the hydrated films was measured by Fluorescence Recovery After Photobleaching (FRAP). Hydrated films cut into squares of ca. 1 cm^2^ were immersed overnight in FITC-dextran solutions (2000 ppm) with molecular weight 10, 40 and 500 kDa. The FRAP measurements were performed using a Zeiss LSM780 confocal microscope. A 488 nm excitation laser was used to bleach a 50 µm diameter circle. The half-time of fluorescence recovery into the circle was then used to characterize the diffusion.

### Strain construction

All plasmids, primers and genetic sequences relevant to this study are listed in **Table S1**. All cloning steps were performed in Escherichia coli XL1-Blue. Golden Gate assembly was used for construction of pPMQAK1-CRISPR/Cas9 target vector by mixing all genetic parts (sgRNA, donor DNA and pPMQAK1-CRISPR/Cas9 base vector) with Thermo Fisher Scientific T4 DNA ligase and FastDigest BsaI, by following step-by-step the plasmid construction protocol from Cengic et al., 2022 (Cengic et al., 2022). NebBuilder HiFi DNA Assembly strategy was used to create integrative and replicative vectors by mixing HiFi Master Mix and PCR products.

The nonmotile glucose-tolerant cyanobacterial strain *Synechocystis* sp. PCC 6803 (Kaplan) was used to develop strain SLS21. Each intermediate and final mutant used in this study are described in **Supplementary Table S2**. A *ggpS sll1566* knock-out was generated by replacing the target gene with an erythromycin resistance (Em) cassette via homologous recombination. The integrative plasmid was constructed on a pEERM backbone. The homologous flanking region were PCR-amplified from *Synechocystis* chromosomal DNA 1000 bp upstream and 1000 bp downstream the integrative site sequence. *Synechocystis* cells were transformed via natural transformation at the logarithmic growth phase with 2 µg plasmid DNA and plated on BG11 agar plate containing 20 µg ml^−1^ Em. Colonies were re-streaked on fresh selective solid BG11 plates until fully segregated clones could be confirmed. In the single Δ*ggpS* mutants, invertase gene *sll0626* was inactivated marker less via CRISPR/Cas9 system (Cengic et al., 2022). A self-replicative pDF-lac2 plasmid for the overexpression of the heterologous sucrose permease gene (*CscB* from *E. coli*) and the native sucrose phosphate synthase gene (Sps, sll0045) was kindly provided by (Thiel et al., 2019) and modified as follows. First, spectinomycin resistance cassette was remove, leaving only chloramphenicol resistance gene as selection marker on on the plasmid. The inducible isopropyl β-D-1-thiogalactopyranoside (IPTG) promoter was replaced by the native *Pcpc560* promoter (Zhou et al., 2014). The final RSF1010-derived pDF-lac2 plasmid, named pL12, was transformed into the double mutant (Δ*ggpS*, Δ*inv*) strain by triparental mating with *E. coli* HB101 helper cells with the plasmid pRL443-AmpR as described (Miao et al., 2017) and plated on selective 18 µg ml^−1^ chloramphenicol BG11 plates. The full segregation of the *inv* and *ggpS* genes deletions and the presence of replicative plasmid were verified by amplification of DNA by a colony PCR using a small amount of cell material as template, DreamTaq 2X Green PCR Master Mix (ThermoScientific).

### Incubation and growth of cells

*Synecocystis* stock cultures were grown in liquid BG11 medium buffered with HEPES (20 mM, pH 7.8), at 30°C, under continuous white-light (50 μmol photons m^−2^ s^−1^), in ambient air or enriched with 1% (v/v) CO_2_, if not stated otherwise. Liquid cultures were grown in flat base uncoated 6-well plates (Sarstedt) or Erlenmeyer flasks under constant shaking (160 rpm). Strains maintenance was carried on solid BG-11 plates containing additional 1% (w/v) Bactoagar (brand) and 0.3% (w/v) sodium thiosulphate. For SLS21 cultivation, BG11 medium was supplemented with antibiotics: 20 µg ml^−1^ erythromycin (Em), 18 µg ml^−1^ chloramphenicol (Cm). XL1-Blue *E. coli* cell line was used for all the cloning/ as a generic host for plasmid propagation/ and cultivated in Luria–Bertani (LB) medium at 37°C in a shaker at 150-200 rpm or on the solid LB plates containing 1.5% (w/v) agar, supplemented with appropriate antibiotics (20 µg ml^−1^ Em and 34 µg ml^−1^ Cm).

### Morphological and spectroscopic characterisation of the immobilised photosynthetic cells

The morphology of the immobilized photosynthetic colonies (wild type, 30°C and 3 µE) was characterized using a laser scanning confocal microscope (LSM780, Carl Zeiss) equipped with a 20X/0.8 NA objective. Chlorophyll autofluorescence was excited at 633 nm and detected using a 650 nm long-pass filter. The detection pinhole was set to 1 Airy unit, yielding a theoretical axial resolution of 1.39 µm. Volumetric image stacks were acquired over a 212.5 µm × 212.5 µm field of view with an axial step size of 0.4 µm, oriented perpendicular to the CNF biohybrid cross-section. This sampling interval fulfills the Nyquist criterion for the established axial resolution. Image processing and quantification were performed using ImageJ (Schneider et al., 2012). Following an intensity-based thresholding of the autofluorescence signal, average cell dimensions were calculated using the 3D Particle Analyser plugin. Chlorophyll *a* (Chl) concentration of the immobilised cells was determined by incubating each biohybrid disc (diameter 1.78 cm; height 0.02 cm) in 3 mL of 90% (v/v) methanol. For liquid cultures, 1 mL of culture was centrifuged, and the resulting pellet was resuspended in 1 mL of 90% methanol. All samples were incubated overnight at 4°C in the dark. After incubation, samples were centrifuged at 4°C at 4800 rpm for 5 min, and the absorbance of the supernatant was measured at 665 nm using a UV-1800 spectrophotometer (Shimadzu, Japan). Chlorophyll *a* concentration was calculated according to Lichtenthaler using an extinction coefficient of 12.9 mg L⁻¹ cm⁻¹ ( Lichtenthaler, 1987).

### Sucrose production in liquid and immobilised cultures

Suspension cultures for sucrose production were grown in BG11 medium buffered with HEPES (20 mM, pH 7.8) and supplemented with appropriate antibiotics. Cultures were maintained in a growth chamber with orbital shaker (160 rpm), at 30°C, either under 50 μmol photons m^−2^ s^−1^ and enriched air with 1% (v/v) CO_2_ or under 3, 15 μmol photons m^−2^ s^−1^ in ambient air. Pre-cultures were grown until reaching OD_730_: 1.5, after which cells were pelleted and resuspended to a final OD_730_: 0.3 in BG11 containing 0.2 M NaCl for salt adaptation. After 24 h, cultures were centrifuged again and resuspended to OD_730_: 0.3 in BG11 supplemented with 0.4 M NaCl, as it shown to be the optimal concentration for induced sucrose production (Tóth et al., 2022). Cultures were grown either in 100 mL Erlenmeyer flasks or in flat-bottom uncoated 6-well plates. Stock cultures intended for CNF immobilization were treated identically up to the salt-adapted step. Following 24 h in BG11 with 0.2 M NaCl, cells were pelleted and resuspended in BG11 without added salt before mixing with CNF. The resulting CNF-immobilized cell discs were cultivated in 4 ml BG11 containing 0.4 M NaCl in uncoated 6-well plates. Immobilised cells were maintained in a semi-continuous mode: at each sampling point for sucrose analysis, the culture medium was fully replaced with fresh BG11 containing 0.4 M NaCl. For sucrose quantification, aliquots from suspension cultures and from the media surrounding CNF-immobilized cells were centrifuged at maximum speed for 10 min at 4 °C. Supernatants were either analyzed immediately or stored at - 20 °C until further processing. Sucrose was quantified via HPLC. The chromatographic separation of sodium chloride, sucrose, fructose and glucose was performed on Vanquish-HPLC (ThermoFisher) combined with VH-A10-A automatic sampler and Vanquish binary pump VH-P10-A. The system is equipped with a Hi-Plex H USP column L17 (7.7 × 100 mm, 8 μm) (Agilent Technologies, Inc., UK), kept at 45°C, and Vanquish™ Refractive Index Detector (RID) (Thermo Scientific™). The mobile phase solution was 0.005 N H_2_SO_4_ (20 mM sulfuric acid) at a flow rate of 0.4 ml min^-1^ and it was also used to flush the syringe of the auto sampler. The injected volume was 10 μl and the total run was completed in 15 min (Zaky et al., 2017).Calibration curves for sucrose, fructose and glucose in 0-5 mM range were run together with samples.

### Proteomic analysis by LC-MS/MS and sample preparation

Liquid and immobilised SLS21 cultures were collected at day 3, 6 and 9 of experiment and treated to obtain cell extract for proteomic analysis. CNF discs were frozen at -20°C directly at the collection time, while 1 mL of liquid culture was pelleted by centrifuging at 3000 rpm, 5 min at RT and stored at -20°C. Sample buffer (100 mM PBS in pH 7-8.5) containing protease inhibitors (cOmplete, EDTA-free Protease Inhibitor Cocktail, Roche) was used to resuspend the cells pellet and immobilised cells in 250 μl. The samples were mixed with 100 μL glass beads (diameter 425-600 μm, Sigma-Aldrich) in a FastPrep lysis tube and lysed by bead beating (FastPrep-24 5G lysis machine, MP Biomedicals) over six cycles of 45 s at 6.5 m s^−1^ and 4°C, with 30 s breaks on ice between cycles. The lysed cells were then pelleted by centrifugation (13,000 rpm, 5 min, 4°C) and supernatants were transferred to new tubes. Lysate protein concentration was determined using a Bradford assay using BSA as a standard, after which samples were diluted to a concentration of 1 μg μL^-1^. For some samples, lysates were too dilute, in which case a larger volume of peptide digest was used for C18 desalting to compensate. Subsequently, samples were reduced in mM DTT for 45 min at 55°C and alkylated in 17 mM IAA for 30 min at RT. Proteins were digested for 18 hours with Trypsin/LysC Mix at a 1:50 protease to protein ratio, after which the reaction was quenched by addition of formic acid, reducing sample pH to below 3. Samples were desalted using stage tips packed with 6 layers of C18 Empore SPE disks with the following chromatographical workflow: activation with 50 μL acetonitrile, equilibration with 200 μL 0.1% formic acid, sample application, 2x wash with 200 μL 0.1% formic acid and 2x elution with 30 μL 80% acetonitrile, 0.1% formic acid. After application in all steps, the stage tips were centrifuged at 3000 rpm until all liquid had passed through the tips. Samples were then evaporated to dryness at 40°C in a speedvac before being resuspended in 20 μL 0.1 % formic acid and stored at -20 °C until mass spectrometry analysis. 12.5 μg of each sample was resuspended in 20 μL 20 mM EPPS buffer pH 8.5 and labelled with 0.1 mg TMTpro 18plex isobaric reagent for one hour, before quenching with 5% hydroxylamine and pooling each set of 18 labelled samples. The pooled samples were then fractionated into 6 fractions using Pierce High pH Reversed-Phase Peptide Fractionation Kit and dried with vacuum centrifugation at 45°C. Dry samples were stored at -20°C before being resuspended in 15 μL 0.1% formic acid prior to LC-MS/MS injection. The samples were analyzed using a Q-Exactive HF Hybrid Quadrupole-Orbitrap mass specgtrometer coupled to an UltiMate 3000 RSLCnano system with an EASY-Spray ESI ion source. 2 μL sample was loaded onto a C18 Acclaim PepMap 100 trap column (75 μm × 1.5 cm, 3 μm, 100 Å) with a flow rate of 7 μL min^-1^, using 3% acetonitrile, 0.1% formic acid and 96.9% water as solvent. The samples were then separated on ES902 EASY-Spray PepMap RSLC C18 Column (75 μm × 25 cm, 2 μm, 100 Å) with a flow rate of 0.7 μL min^-1^ using a 120 min long linear gradient from 1% to 32% with 95% acetonitrile, 0.1% formic acid and 4.9% water as secondary solvent. Data was acquired using a single full scan (resolution 120,000 at 200 m/z, mass range 350-1500 m/z) followed by 15 MS2 DDA scans using the 15 most abundant peptides. For MS2 scans, a resolution of 120,000 at 200 m/z was maintained, with an isolation window of 0.7 m/z. Precursor ions were fragmented using high-energy collision-induced dissociation at an NCE of 30. Maximum injection time and automatic gain control were set to 50 ms and 3E6 for MS1, and 250 ms and 1E6 for MS2, respectively. Raw data were processed with Thermo Fisher Proteome Discoverer v. 3.259 using the SEQUEST engine. Normalized peptide intensities were exported for further processing. A FASTA file from the UniProt proteome UP00000142561 was used for the search with relevant heterologous proteins added in manually. Oxidation (M) and Acetyl (Protein N-term) were used as variable modifications and Carbamidomethyl (C) as a fixed modification. A maximum of two missed cleavages were allowed, and the false discovery rate was set to 1%. Samples were median normalized and fold-changes were calculated using the MSstats (v. 4.4.1) R package (Choi et al., 2014). The p-values were adjusted for multiple hypothesis testing using the Benjamini-Hochberg method, with an adjusted p-value threshold of 0.05. Proteins detected in fewer than 3 replicates were excluded from statistical analysis.

## Author contributions

Conceptualization: SL, EPH, VG

Methodology: SL, ES, EPH, VG

Investigation: SL, ES, NLA, VA, EK, HB, TM, EPH, VG

Visualization: SL, ES, VG

Supervision: EPH, VG

Writing-original draft: SL, ES, NLA, VA, EPH, VG

Writing-review & editing: SL, TM, EK, HJ, EPH, VG

